# Assessment of *Plasmodium falciparum* infection and fitness of genetically modified *Anopheles gambiae* aimed at mosquito population replacement

**DOI:** 10.1101/2021.11.01.466738

**Authors:** Sofia Tapanelli, Maria Grazia Inghilterra, Julia Cai, James Philpott, Paolo Capriotti, Nikolai Windbichler, George K. Christophides

## Abstract

Genetically modified (GM) mosquitoes expressing anti-plasmodial effectors propagating through wild mosquito populations by means of gene drive is a promising tool to support current malaria control strategies. The process of generating GM mosquitoes involves genetic transformation of mosquitoes from a laboratory colony and, often, interbreeding with other GM lines to cross in auxiliary traits. These mosquito colonies and GM lines thus often have different genetic backgrounds and GM lines are invariably highly inbred, which in conjunction with their independent rearing in the laboratory may translate to differences in their susceptibility to malaria parasite infection and life history traits. Here, we show that laboratory *Anopheles gambiae* colonies and GM lines expressing Cas9 and Cre recombinase vary greatly in their susceptibility to *Plasmodium falciparum* NF54 infection. Therefore, the choice of mosquitoes to be used as a reference when conducting infection or life history trait assays requires careful consideration. To address these issues, we established an experimental pipeline involving genetic crosses and genotyping of mosquitoes reared in shared containers throughout their lifecycle. We used this protocol to examine whether GM lines expressing the antimicrobial peptide (AMP) Scorpine in the mosquito midgut interfere with parasite infection and mosquito survival. We demonstrate that Scorpine expression in the Peritrophin 1 (Aper1) genomic locus reduces both *P*. *falciparum* sporozoite prevalence and mosquito lifespan; both these phenotypes are likely to be associated with the disturbance of the midgut microbiota homeostasis. These data lead us to conclude that the Aper1-Sco GM line could be used in proof-of-concept experiments aimed at mosquito population replacement, although the impact of its reduced fitness on the spread of the transgene through wild populations requires further investigation.

## Introduction

Engineering the mosquito genome has become a key resource in renewed efforts to control the spread of malaria (Hammond and Galizi, 2017). Following the establishment of a reliable genome transformation method in *Anopheles* mosquitoes (Catteruccia et al., 2000;Grossman et al., 2001), several studies explored the use of homing endonucleases (HEGs), including Cas9, in conjunction with gene drive as a method to spread engineered genetic traits in wild vector populations (Windbichler et al., 2011;Hammond et al., 2016;Kyrou et al., 2018;Carballar-Lejarazu et al., 2020). Such traits are aimed at either reducing the vector effective population size (population suppression) or rendering the vector less capable of transmitting malaria (population replacement).

We have previously established a novel approach, termed Integral Gene Drive (IGD), which allows us to introduce minimal genetic modifications in any *A*. *gambiae* gene of choice making it able to express small effector proteins targeting the malaria parasite, including exogenous AMPs (Nash et al., 2019;Hoermann et al., 2021). Creation of an IGD effector line entails precision engineering of the endogenous gene of choice using CRISPR/Cas9, by introducing a sequence that encodes the small effector protein in frame with the endogenous protein-coding sequence; the two sequences can be separated with the insertion of a 2A ribosomal skipping sequence or remain fused. The exogenous sequence contains two additional expression cassettes often found within an intron: the first produces the guide RNA (gRNA) that allows the engineered allele to home in the presence of Cas9, and the second expresses a fluorescent reporter gene that serves as a transformation marker, flanked by LoxP sites that enables it to be removed at a later stage in the presence of Cre recombinase.

*P*. *falciparum*, the deadliest of human malaria parasites, spends about a day inside the mosquito midgut after being ingested with a bloodmeal and before it establishes an infection in the midgut wall in the form of an oocyst (Sinden, 2002). In our first proof-of-concept of the IGD protocol, we expressed the AMP Scorpine in the adult female *A*. *gambiae* midgut, targeting early malaria transmission stages (Hoermann et al., 2021). This was achieved by introducing Scorpine in either of three bloodmeal-induced loci expressing the blood-processing enzymes carboxypeptidase (CP) and alkaline phosphatase 2 (AP2) and the type I peritrophic matrix protein Peritrophin 1 (Aper1). While Scorpine was separated from CP and AP2 through a 2A ribosomal skipping peptide, it was designed to remain fused with Aper1, leading to it being anchored to the peritrophic matrix. The three resulting lines were designated as Sco-CP, ScoG-AP2 and Aper1-Sco, respectively. The generation of these three lines required genetically transformed mosquitoes of the *A*. *gambiae* G3 laboratory colony to be crossed to a KIL line expressing Cre-recombinase (KIL^vasa-Cre^) to remove the fluorescent marker and a G3 Cas9-expressing line (G3^v*asa*-Cas9^) to facilitate homing and homozygosity of the transgene.

The next step in the path toward establishing whether an IGD line displays the qualities required for a population replacement strategy involves a series of laboratory assays to assess malaria transmission reduction efficacy and any potential impact of the modifications on mosquito life history traits (James et al., 2020). Assessment of malaria transmission reduction typically involves enumeration of oocysts in the mosquito midguts following infection with *P*. *falciparum* gametocytes using a standard membrane feeding assay (SMFA) (Habtewold et al., 2019). Infection prevalence and intensity are compared to a reference line, typically the parental line where the modifications have been performed. Using this protocol, we showed that two of the Scorpine-expressing lines, the ScoG-AP2 and the Aper1-Sco, exhibit significant reduction of the number of *P*. *falciparum* oocysts found in the mosquito midgut following an infectious bloodmeal compared to the G3 parental line.

Recent studies have established that the protocol mentioned above may not provide the most reliable measure of transmission reduction as it does not consider the oocyst developmental stage, growth rate and capacity to produce sporozoites (Shaw et al., 2020;Habtewold et al., 2021). Furthermore, these studies suggest that provision of regular supplemental bloodmeals following the initial infectious bloodmeal, each followed by egg laying, must be considered, as this better emulates the mosquito feeding and reproductive behavior in nature, while they significantly support and, indeed, accelerate oocyst development. Finally, since the reference line (including the parental line) is often genetically different and/or more diverse than the GM effector line to be tested, and is cultured independently of the GM line, the choice of reference requires careful consideration. Many of the above considerations regarding also pertain to the assessment of survival and other life history traits.

Here, we investigate differences in *P*. *falciparum* infection between *A*. *gambiae* laboratory colonies and accessory GM lines used for further genome manipulation, such as G3^v*asa*-Cas9^ and KIL^vasa-Cre^, and establish a new experimental protocol that addresses the aforementioned considerations. Using this protocol, we examine the malaria transmission blocking efficacy of the IGD Scorpine-expressing lines (Hoermann et al., 2021), by assessing the presence and abundance of sporozoites in the mosquito salivary glands. Finally, we investigate the effect of Scorpine expression within the Aper1 genomic locus on mosquito survival. Our work is fundamental to establishing the most appropriate protocols for testing the characteristics of GM *A*. *gambiae* lines prepared for gene drive and vector population replacement.

## Materials and methods

### Mosquito strains and rearing

The *A. coluzzii* N’gousso colony was established from field-collected mosquitoes of the *A*. *gambiae* M molecular form in 2006 in Cameroon (Harris et al., 2010). The *A*. *gambiae* G3 strain was obtained from MR4 and is classified as *A*. *gambiae* S molecular form. The *A*. *gambiae* G3 vasa-Cas9 (G3^vasa-cas9^), KIL vasa-Cre (KIL^vasa-Cre^) and effector IGD (ScoG-AP2, Sco-CP and Aper1-Sco) strains were a kind gift of their respective creators (Volohonsky et al., 2015;Kyrou et al., 2018;Hoermann et al., 2021). All strains were reared and maintained at standard laboratory conditions (27 ± 1 °C and 70 ± 5%) humidity on a 12:12 light/dark cycle interspersed by 0.5 h dawn and dusk transitions.

### *P*. *falciparum* culturing and SMFA

*P*. *falciparum* NF54 strain was obtained from the MR4 (MRA-1000, patient E). Parasite culturing and mosquito SMFAs were performed as described (Habtewold et al., 2019). Briefly, asexual parasite stages were cultured in RPMI-1640-R5886 (Sigma) medium supplemented with hypoxanthine to end concentration of 0.05 g/L, L-glutamine powder-(G8540-25G Sigma) to end concentration of 0.3 mg/L and 10% sterile human serum. Gametocytogenesis was induced by diluting the asexual culture (3-4% ring forms) to 1% ring forms with fresh human red blood cells (hRBCs) and maintained at 37°C, ensuring a daily exchange of about 75% of the medium. Day 14-16 mature gametocytes were pooled in a pre-warmed 15 mL tube containing 20% hRBCs and 50% heat-inactivated human serum. Each membrane feeder was loaded with 300 μl of the gametocyte mixture and kept at a constant temperature of 37°C. Paper cups of 70-100 female mosquitoes were allowed to feed for 10-15 minutes. After the feed, mosquitoes were maintained in a chamber set at 27°C and 70% humidity. For oocyst enumeration, mosquito midguts were dissected 7 days post-bloodmeal, and for sporozoite quantification, mosquitoes were dissected at day 14 post-infection, after completing 4 full gonotrophic cycles including supplementation with non-infectious bloodmeals at days 4, 6 and 9 and egg laying at days 3, 5, 8 and 11 post-infection. For oocyst enumeration, mosquito midguts were stained with 1% mercurochrome and fixed in 4% formaldehyde, and oocysts were counted using a bright-field light microscope at 20X magnification. For sporozoite quantification, mosquito heads and thoraxes were separated from the abdomens using a scalpel and preserved in TRIzol reagent (Invitrogen) at −80°C until processed for RNA extraction. The abdominal section of individual mosquitoes was used for genotyping PCR.

### Mosquito genotyping

Individual mosquitoes were genotyped using Phire Tissue Direct PCR Master Mix (Thermo-Scientific). PCR was performed according to the manufacturer’s protocol using genomic DNA extracted from mosquito abdomens as template. The PCR primers used for the genotyping assays were: CP-locus-F: ATTATCGGAAATTCTCCACAAAGCG, CP-locus-R: TCATGGTGCGATTCTGATGCGATCG, AP-locus-F: ATCAGACTTTAAGCCTTTGTCGTC and AP-locus-R: ACCGTTCGCGATTCAGCAGTACAGG.

### Quantification of *P*. *falciparum* sporozoites

RNA from mosquito heads and thoraxes was extracted using the TRIzol™ reagent according to the manufacture’s guidelines (Invitrogen). RNA samples were stored at −80°C until cDNA synthesis using the iScript™ gDNA Clear cDNA Synthesis Kit according to the manufacture’s guidelines (BIO-RAD). A qPCR-based assay was used to detect and quantify *PfCSP* transcripts (Kefi et al., 2018), normalized against the *A*. *gambiae* ribosomal *S7* gene (Dimopoulos et al., 1998). The qPCR was performed using SYBR™ Green PCR Master Mix (Thermo Fisher Scientific) using primers: *PfCSP*-F: TCAACTGAATGGTCCCCATGT, *PfCSP*-R: GAGCCAGGCTTTATTCTAACTTGAAT, S7-F: GAGGTCGAGTTCAACAACAA, and S7-R: GAACACGACGTGCTTACC. Samples were considered not infected when no Cp value was calculated using Absolute quantification/2^nd^ derivative Max analysis on the Light Cycler® 480 (Roche, Software release 1.5.1.62 SPS).

### Survival assay

Mosquitoes were provided supplemental bloodmeals every three days and allowed mosquitoes to deposit eggs. After the first blood meal, individual dead mosquitoes were collected every 24 hours until the caged population was extinct. Mosquito carcasses were used for genotyping.

### Microbiota analysis

DNA was extracted from whole mosquitoes collected 24 hours after the first or second bloodmeal and DNA. The total load of bacteria and bacterial families were measured on the 7500 Fast Real-Time PCR system (Applied Biosystems) in 10 μL reactions using Fast SYBR™ Green PCR Master Mix (Thermo Fisher Scientific) and *16S* primers: 16S-F: TCCTACGGGAGGCAGCAGT, 16S-R: GGACTACCAGGGTATCTAATCCTGTT, Enterobacteriaceae 16S-F: CGTGCTACAATGGCATATACAAAGAGAAG, Enterobacteriaceae 16S-R: AGCATTCTGATCTACGATTACTAGCGATTC, Flavobacteriaceae 16S-F: TAAGGTTGAAGTGGCTGGAATAA, and Flavobacteriaceae 16S-R: GTCCATCAGCGTCAGTTAAGACT. Expression of 16S was normalized against S7.

### Statistical analyses

Infection intensity was assessed by the enumeration of oocysts and by molecular quantification of sporozoites using qPCR. We used Mann-Whitney test for non-normal, unpaired data sets. The prevalence of infections and the difference in the survival percentage of mosquitoes was evaluated with the Chi-squared Pearson’s test. Estimates of the mosquito survival probability was performed with the Kaplan-Meier statistics. All statistical analyses were preformed using R (version 3.5.3) and Graph Pad Prism version 8.0.

## Results

### Laboratory *A*. *gambiae* susceptibility to *P*. *falciparum* infection

We previously established a robust protocol for SMFAs using the *A*. *coluzzii* laboratory colony N’gousso (Habtewold et al., 2019). *A*. *coluzzi*, until recently known as M-form *A*. *gambiae*, is the main malaria vector in West sub-Saharan Africa (della Torre et al., 2005). However, this species is not present in East Africa, where one of the main vectors is *A. gambiae sensu stricto (ss)*, previously known as S-form *A*. *gambiae* and henceforth simply referred to as *A*. *gambiae*. Therefore, we had used *A*. *gambiae* G3 strain, the most widely and long-term used *A*. *gambiae* laboratory colony, for generating GM mosquitoes.

We compared the *A*. *gambiae* G3 (henceforth G3) and *A*. *coluzzii* N’gousso (henceforth N’gousso) with regards to their *P*. *falciparum* infection intensity and prevalence using data obtained from 19 and 16 independent, non-paired infection experiments, respectively (<u>Figure 1A</u>). The results showed that G3 exhibits a significantly lower infection intensity (p<0.0001, unpaired two-tailed Mann-Whitney test) with an oocyst mean of 11 (SD: +/−29.95, SEM: +/−1.38), median of 0 and range of 0-348 compared to N’gousso with an oocyst mean of 35 (SD: +/−70.26, SEM: +/−2.7), a median of 4 and range of 0-486 oocysts per midgut, respectively. The infection prevalence was also lower (p<0.001, Chi-squared Pearson’s test) in G3 (67%) compared to N’gousso (43%).

**Figure 1.**
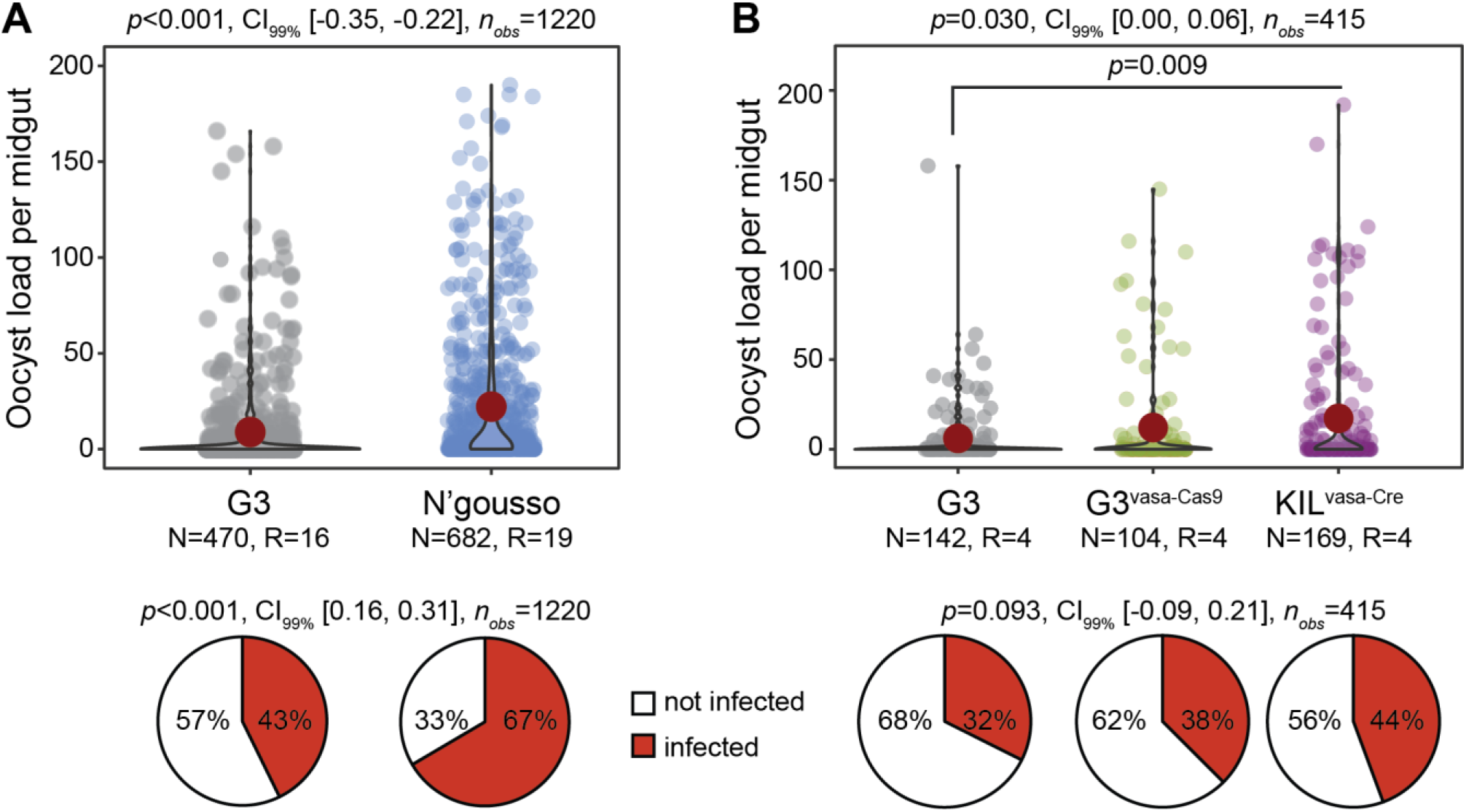
Susceptibility of laboratory *A*. *gambiae* lines to *P*. *falciparum* NF54. **A**. Infection intensity and prevalence (%) of G3 and N’gousso. **B**. Infection intensity and prevalence of G3, G3^vasa-Cas9^ and KIL^vasa-Cre^ shown as a pool of four biological replicates. The violin dot plots show the frequency and distribution of data. Single dots show the number of oocysts counted per midgut and the red circle shows the median infection intensity. N is the number of mosquitoes analyzed per line and R is the number of biological replicates. Statistical significance is shown on top of the graphs (*p*), with the CI_99%_ and the number of observations analyzed per experiment. The pie charts show the percentage of not infected (white) and infected (red) mosquitoes.

To generate GM mosquito effector lines using the IGD design (Nash et al., 2019), GM G3 mosquitoes had to be crossed to a line that expresses Cre recombinase (here KIL^vasa-Cre^; (Volohonsky et al., 2015) to remove the transformation marker that is flanked by LoxP sites (Hoermann et al., 2021) and another line expressing Cas9 (here G3^vasa-Cas9^; (Volohonsky et al., 2015) to achieve homozygosity through homing of the transgene (Hoermann et al., 2021).

Therefore, a homozygous IGD effector lines has mixed genetic backgrounds. We examined any potential differences in *P*. *falciparum* infection between the G3, G3^vasa-cas9^ and KIL^vasa-Cre^ lines that were used to generate the IGD effector lines Sco-Cp, ScoG-AP2 and Aper1-Sco (Hoermann et al., 2021). Results from four independent infections revealed no difference between the parental G3 and the G3^vasa-cas9^ lines. However, both lines were significantly different from the KIL^vasa-Cre^ strain that showed higher infection intensity (<u>Figure 1B</u>; p=0.03, two-tailed unpaired Mann-Whitney test). These data suggest that neither the G3 nor any of the G3^vasa-cas9^ and KIL^vasa-Cre^ lines serve as good references for the GM effector line.

### Assessment of infection blocking efficacy of GM effector lines

Two factors were thought to be responsible for the infection variability detected between the various colonies and lines: the mosquito diverse genetic background and enterotype. Whilst the former can be addressed in many different ways (*e.g.*, backcrossing the effector line to the parental G3 colony and comparing the progeny with the G3 control or establishing a control population through crosses of the various founder lines) the latter can only be effectively addressed by rearing mosquitoes in the same containers including the aquatic larval stages. We designed an assay to concurrently address these issues, whereby the parental GM effector line that had a mixture of G3 and KIL genetic backgrounds was backcrossed to the G3 line (referred to here as wild type, wt, with regards to the integration locus) to obtain hemizygote transgenic mosquitoes of generation G_1_ which were then allowed to interbreed to derive a generation G_2_ of mosquitoes (<u>Figure 2A</u>). This new isogenic population of mosquitoes is comprised of wt homozygotes (−/−), transgenic hemizygotes (+/−) and transgenic homozygotes (+/+). As all mosquitoes are reared together, they are expected to be exposed to the same microbiota and thus share the same enterotype.

**Figure 2.**
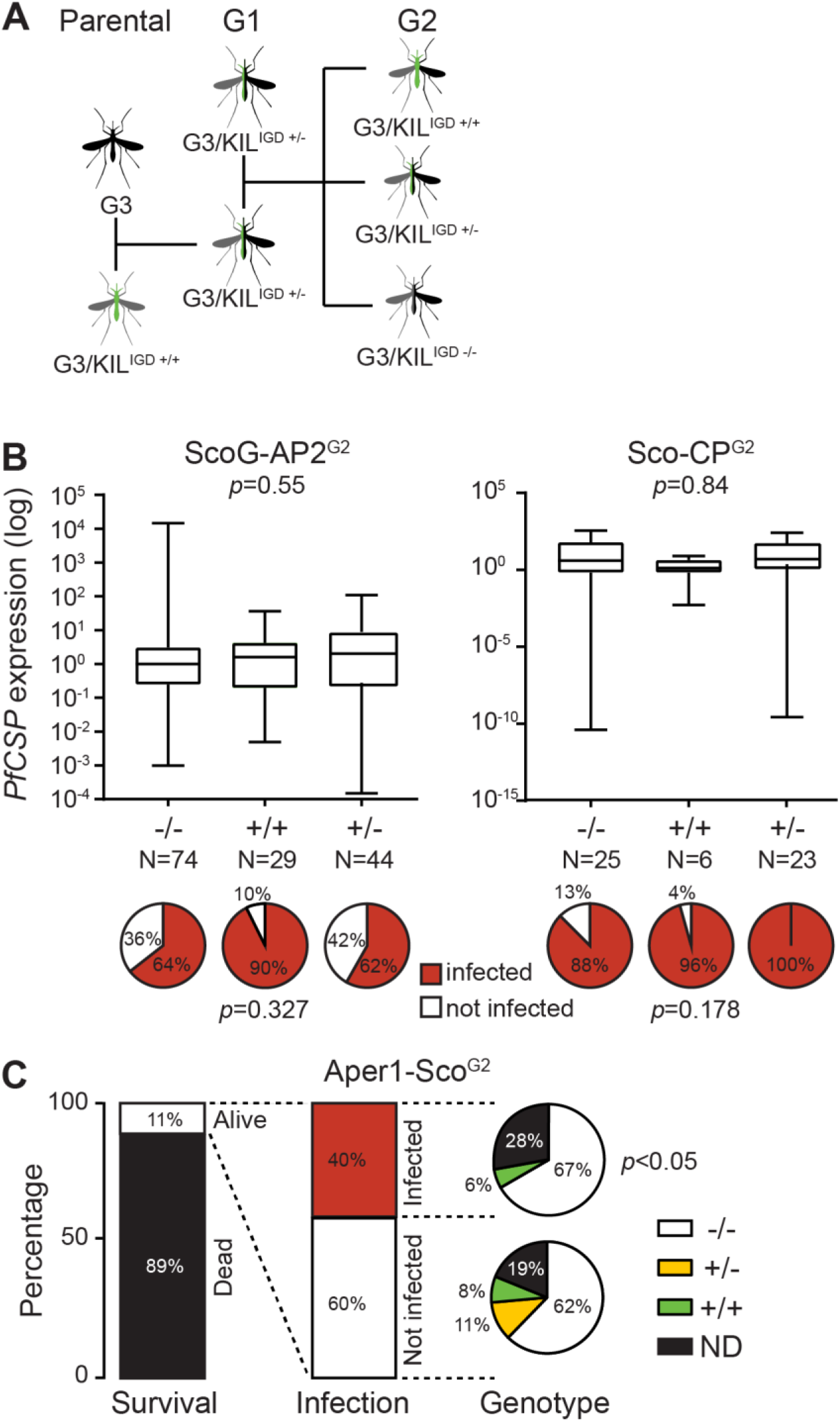
*P*. *falciparum* NF54 infection of Scorpine-expressing GM *A*. *gambiae* lines. **A.** Schematic representation of crosses required for the analysis of the IGD effector lines. **B.** Quantification of sporozoites in ScoG-AP2^G2^ (left) and Sco-CP^G2^ (right) mosquitoes by qPCR of *PfCSP* transcripts in preparations of mosquito heads and thoraxes, normalized on housekeeping AgS7 and analyzed using the Mann-Whitney test. Pie charts below each graph show the prevalence of infection (sporozoite presence; %) in each sub-population analyzed using a Chi-Squared Pearson’s test. N is the number of mosquitoes dissected and analyzed. **C.** Histograms report the percent survival and *P*. *falciparum* infection of the Aper1-Sco^G2^ population 14 days post-infectious bloodmeal, respectively; the latter determined by qPCR of *PfCSP* transcripts. Pie charts report the genotype composition of infected and non-infected mosquitoes, analyzed using a Chi-Squared Pearson’s test.

We used this assay to test three IGD GM effector lines we previously generated, which express the AMP Scorpine in three different *A*. *gambiae* genomic loci: Sco-Cp, ScoG-AP2 and Aper1-Sco (Hoermann et al., 2021). Female mosquitoes of each population were infected 3 days post-emergence with *P*. *falciparum* gametocytes provided through a SMFA. To further emulate the mosquito natural behavior and physiology, mosquitoes were provided 3 supplemental non-infectious bloodmeals (day 3, 6 and 9 post-infection) through membrane feeding, each followed by egg laying. At day 14 post-infection, surviving female mosquitoes were dissected to molecularly quantify salivary gland *P*. *falciparum* sporozoites by assessing the presence and abundance of Circumsporozoite protein (PfCSP) transcripts in mosquito heads and thoraxes using quantitative real-time PCR (qRT-PCR) and be genotyped for transgene zygosity by PCR on DNA isolated from their abdomens.

Of 400 ScoG-AP2, 300 Sco-CP and 700 Aper1-Sco mosquitoes pooled from 3 independent replicates each, 240, 60 and 88, respectively, survived to day 14 (<u>Table 1</u>). In all cases, +/+ and +/− transgene genotypes were less than what one would expect from an unbiased Mendelian segregation of alleles, which in conjunction with the high mortality especially in the Sco-CP and Aper1-Sco lines indicated that Scorpine expression may inflict a fitness cost to female GM mosquitoes.

**Table 1.** Genotypes of transgenic lines from heterozygous crosses.

| Line (total number) | Alive [number (% of alive - ND)] |  |  |  | Dead [number] |
| --- | --- | --- | --- | --- | --- |
| Zygosity | -/- | +/- | +/+ | ND |  |
| ScoG-AP2 (400) | 109 (50%) | 62 (30%) | 32 (20%) | 37 | 160 |
| Sco-CP (300) | 23 (47%) | 20 (41%) | 6 (12%) | 11 | 240 |
| Aper1-Sco (700) | 55 (81%) | 6 (9%) | 7 (10%) | 20 | 612 |

No significant differences were observed between +/+, +/− and −/− ScoG-AP2 and Sco-CP, respectively, both with regards to *PfCSP* transcript abundance (Mann-Whitney test, p=0.55 and p=0.84 respectively) and presence (Chi-Squared Pearson’s test, p=0.32 and p=0.17, respectively) (<u>Figure 2B</u>). However, of the 88 Aper1-Sco female mosquitoes that survived to day 14, 35 (40%) were found to have PfCSP transcripts, of which 23 were −/−, 2 were +/+ and 0 were +/−, revealing a significant difference in sporozoite prevalence between transgenic (both +/+ and +/−) and non-transgenic mosquitoes (<u>Figure 2C</u>; Chi-squared Pearson’s test, p<0.05). We could not determine the genotype of the remaining 10 mosquitoes. The very low numbers of sporozoite-positive transgenic mosquitoes did not permit any meaningful statistical analysis of *PfCSP* transcript abundance.

### Aper1-Sco expression reduces the survival probability of aged mosquitoes

To gain further insights into the survival of Aper1-Sco-expressing mosquitoes, we used the same assay but now genotyped dead female mosquitoes of generation G_2_ (Aper-Sco^G2^) every 24 hours starting from day 1 after the first (here non-infectious) bloodmeal. The overall survival probability of the Aper-Sco^G2^ female population was also compared to that of the parental G3 female population. All mosquitoes were blood-fed every 3 days and each bloodmeal was followed by egg laying.

Compared to G3, the Aper1-Sco^G2^ female population exhibited significantly lower probability of survival over time (Kaplan-Meier statistics, p<0.0001; <u>Figure 3A</u>). Importantly, only 5% of Aper1-Sco^G2^ female mosquitoes (21 of 429) survived to day 10 post first bloodmeal, when *P*. *falciparum* is known to reach its peak infectivity to humans, compared to 41% of G3 female mosquitoes (143 of 353; Chi-squared Pearson’s test, p<0.0001). While some females of the parental G3 line survived to day 20, the Aper1-Sco^G2^ line was fully suppressed by day 14. Within the Aper1-Sco^G2^ population, the survival probabilities of +/+ and +/− female mosquitoes appeared to be drastically reduced compared to −/− mosquitoes although only +/− and −/− mosquitoes showed significantly different survival probabilities with Kaplan-Meier statistics (p<0.001; <u>Figure 3B</u>). Specifically, only 13% of +/− (Chi-squared Pearson’s test, p<0.001) and 19% of +/+ (Chi-squared Pearson’s test, p<0.01) female mosquitoes survived after day 6 compared to 42% of −/− female mosquitoes.

**Figure 3.**
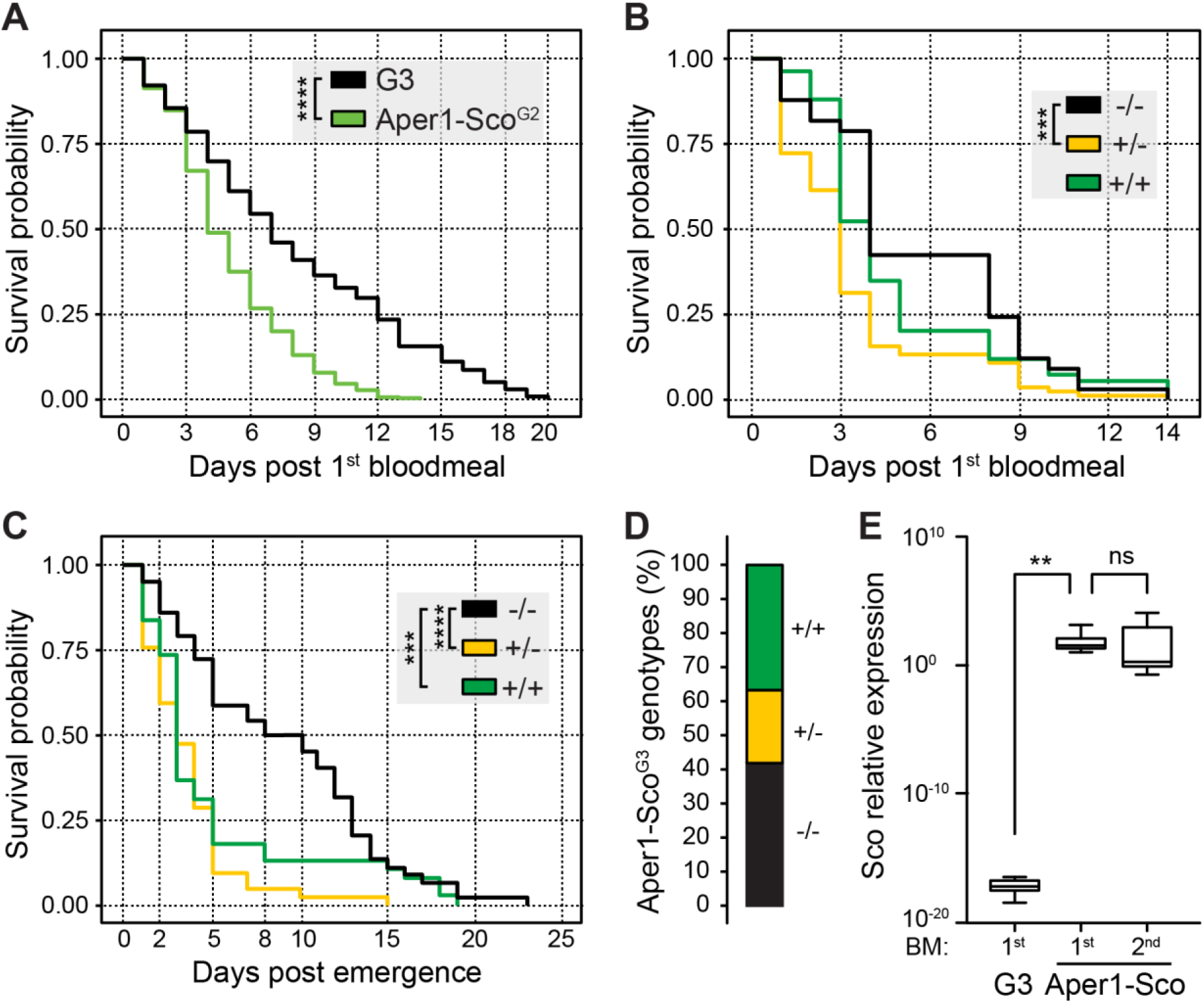
Survival and reproduction assays of the Aper1-Sco^G2^ mosquito population. Kaplan-Meier survival curves and statistical analysis of survival probabilities of G3 and Aper1-Sco^G2^ female mosquito populations (**A**) and each IGD effector genotype (−/−, +/− and +/+) of the Aper1-Sco^G2^ female population (**B**). In both assays, day 0 is the day of the 1^st^ bloodmeal, and supplemental bloodmeals were provided every 3 days thereafter. (**C**) Kaplan-Meier survival curves and statistical analysis of survival probabilities of individual genotypes of the Aper1-Sco^G2^ mosquito population including both females and males. In this assay, day 0 is the day of adult emergence, while the 1^st^ bloodmeal was provided on day 2 and the two additional bloodmeals were provided on days 5 and 8 post-emergence. (**D**) Percentage of genotypes in the adult Aper1-Sco^G3^ population (both females and males) that was derived from eggs laid after each of the bloodmeals of Aper1-Sco^G2^ mosquitoes of panel C. (**E**) Relative expression of Scorpine (Sco) in control G3 and Aper1-Sco +/+ mosquitoes 24 hours after the 1^st^ or 2^nd^ bloodmeal. Expression of Scorpine was normalized against the *A. coluzzii S7* transcripts. The analysis was performed using a student’s t test. ** p<0.01; *** p<0.001; *** p<0.0001.

Next, we investigated the capacity of the Aper1-Sco^G2^ population to produce offspring that inherit the transgene in the absence of gene drive. Following crosses and rearing of mosquitoes in the same containers as described above, the adult Aper1-Sco^G2^ population was fed on non-infected human blood on days 2, 5 and 8 post-adult emergence. After every bloodmeal, females laid eggs and adults that emerged from these eggs (Aper1-Sco^G3^) were pooled and genotyped for the presence of the Aper1-Sco transgene. The parental Aper1-Sco^G2^ population was cultured until all male and female mosquitoes had died and then genotyped. The results showed that the founder Aper1-Sco^G2^ population consisted of 155 adults (66 females and 89 males) of which 45 were −/− (26 and 19 females and males, respectively), 42 were +/− (17 and 25, respectively) and 38 were +/+ (9 and 29, respectively), while the genotype of 30 mosquitoes (14 and 16, respectively) could not be determined. Kaplan Meier survival analysis revealed that both +/− (p<0.0001) and +/+ (p<0.001) Aper1-Sco^G2^ mosquitoes exhibited significantly reduced survival probability compared to −/− mosquitoes, including both females and males (<u>Figure 3C</u>). When female and male survival probabilities were analyzed independently, the results showed that whether +/− or +/+, both female (p<0.0001) and male (p<0.001) mosquitoes died significantly earlier than their wt (−/−) counterparts.

Genotyping of the adult Aper1-Sco^G3^ mosquitoes revealed that all genotypes were represented in the population. Of 111 adults with determined genotypes, 47 (42%) were −/−, 21 (19%) were +/− and 43 (39%) were +/+ (<u>Figure 3D</u>). These data demonstrate that despite their reduced survival probability, largely manifested at later stages of adult life, Aper1-Sco transgenic mosquitoes can produce progeny that inherit the transgene in frequencies resembling those of the wt locus. The bias towards homozygous −/− and +/+ mosquitoes is an interesting observation that deserves further investigation. It is worth noting that owing to the bloodmeal-induced Aper1 locus, Scorpine is over expressed after every bloodmeal, hence its accumulation after several bloodmeals may contribute to the reduced adult mosquito survival (<u>Figure 3E</u>).

### Scorpine expression in the Aper1 locus affects the mosquito gut microbiota

The mosquito gut microbiota community plays an important role in mosquito physiology and interferes with infection with *Plasmodium* (Boissiere et al., 2012;Gendrin et al., 2015;Habtewold et al., 2017;Rodgers et al., 2017;Gao et al., 2020). It is known that the microbiota load and composition are significantly affected by mosquito epithelial immune responses, including AMP expression. To investigate whether Scorpine expression affects the microbiota of the Aper1-Sco mosquitoes, we quantified the bacterial load and recorded the genotype of individual mosquitoes of the Aper1-Sco^G2^ population 24 hours post-bloodmeal by quantitative PCR of the 16S ribosomal DNA. We also separately quantified the relative abundance of Flavobacteriaceae and Enterobacteriaceae, two of the most abundant bacterial families in the *A*. *gambiae* gut (Boissiere et al., 2012;Gendrin et al., 2015;Akorli et al., 2016).

As shown in <u>Figure 4</u>, within the Aper1-Sco^G2^ population we found that mosquitoes of the −/− genotype exhibited a significantly lower abundance of 16S qPCR product compared to +/− and +/+ (Mann-Whitney test, **p<0.01 and *p<0.05, respectively). In the same samples, Enterobacteriaceae were more abundant in +/+ and +/− compared to −/− mosquitoes (Mann-Whitney test, **p<0.01 and *p<0.05, respectively). Since the abundance of Enterobacteriaceae has been associated with reduced *P*. *falciparum* infections (Boissiere et al., 2012;Dennison et al., 2016), these results support the low infection intensity and the statistically lower infection prevalence observed in SMFAs. A similar pattern was evident for the abundance of Flavobacteriaceae, which was higher in both +/+ and +/− compared to −/− mosquitoes (Mann-Whitney, p<0.001, p<0.01 respectively). The increased bacterial load may also be associated with the reduced lifespan of the Aper1-Sco GM mosquitoes (Cansado-Utrilla et al., 2021).

**Figure 4.**
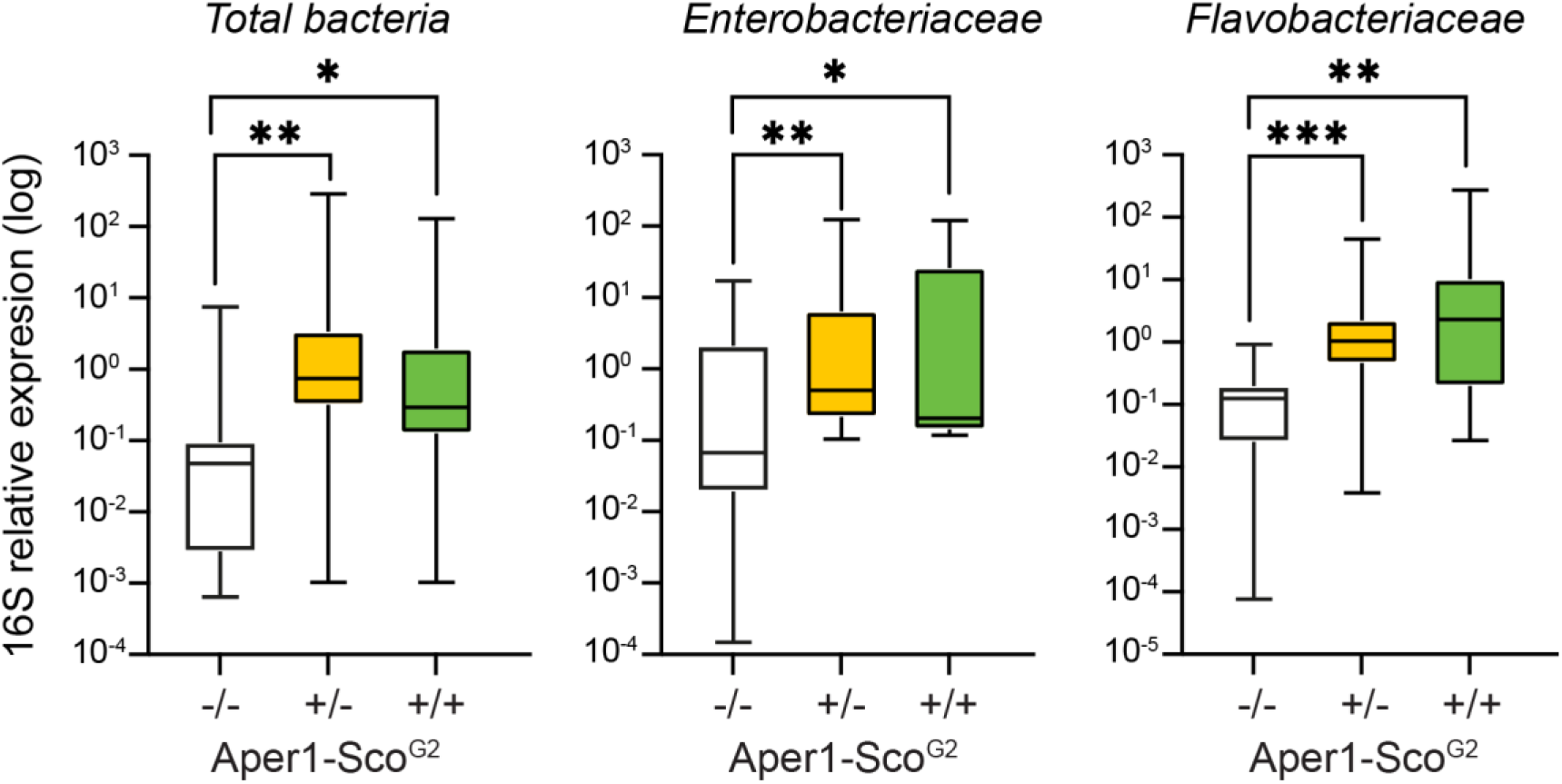
Midgut microbiota load in the Aper1-Sco^G2^ population. Midgut microbiota load in sub-populations of mosquitoes of the Aper1-Sco^G2^ population assessed by 16S qPCR on individual mosquitoes 24 hours post-1^st^ bloodmeal. The loads of total bacteria (left), Enterobacteriaceae (middle) and Flavobacteriaceae (right) were independently assessed. Data from two independent biological replicates were pooled. The numbers of mosquitoes per pool were: for total bacterial load 7 −/−, 29 +/− and 19 +/+; for *Enterobacteriaceae* 10 −/−, 20 +/− and 8 +/+; and for Flavobacteriaceae 8 −/−, 24 +/− and 17+/+. Data were analyzed with a Mann-Whitney test: *, p<0.05; **, p<0.01; ***, p<0.001.

## Discussion

The establishment of robust laboratory assays unambiguously reporting on the various characteristics of a GM *A*. *gambiae* line is a necessary and critical milestone in the path to mosquito population replacement, before large cage, semi-field or field trials can commence. Mosquito vectorial capacity and life history traits can differ substantially between laboratory colonies and lines either because of their inherently different genetic backgrounds or genetic drift caused created through bottlenecks during colonization or establishment of a line. An additional complication is that until recently the incipient species *A*. *gambiae* and *A*. *coluzzii* were interchangeably used and collectively referenced as *A*. *gambiae*, and many publications continue referring to them as such. Moreover, laboratory colonies are often of ambiguous origin or cultured in the same laboratory together with other colonies and lines; since the two incipient species can freely reproduce in the laboratory, contaminations are frequent.

We previously demonstrated that *P*. *falciparum* NF54 infections of the *A*. *coluzzii* N’gousso colony, one of the most recently generated and widely used colonies, produce high infection prevalence and intensity, allowing for the generation of robust data and reliable statistical analyses of transmission blocking (Habtewold et al., 2019). However, this colony cannot be used as a reference for *A*. *gambiae*-derived GM lines, and we show here that N’gousso and G3, the parental colony of GM lines examined here and one of the oldest laboratory colonies, originally of *A*. *gambiae* origin, exhibit markedly different *P*. *falciparum* infection prevalence and intensity, both being significantly lower in G3 than N’gousso. While the G3^vasa-Cas9^ line resembles its parental G3 colony with regards to both infection prevalence and intensity, the KIL^vasa-Cre^ line that is derived from the KIL *A*. *gambiae* strain and has also been used here for the generation of GM effector lines displays significantly higher infection intensity; indeed, KIL was genetically selected to be highly susceptible to infection. To conclude, the choice of reference line must not be done lightly as this may lead to erroneous conclusions.

Whilst there are many ways to choose or generate a reference line that is representative of the genetic variation one expects to have in the GM lines examined, we decided here to develop and use a method that would simultaneously address another fundamental issue in such assays: the culturing of aquatic mosquito stages in different containers. The adult mosquito microbiota are largely representative of what is found in the aquatic larval habitat and are known to affect both *P*. *falciparum* infections and mosquito fitness (Boissiere et al., 2012;Gendrin et al., 2015;Gendrin et al., 2016;Romoli and Gendrin, 2018;Gao et al., 2020). Experience has shown that no matter how similar the conditions and treatment of larval habitats are, the microbiota may differ markedly, even if larval containers are placed side-by-side in the insectary. Therefore, the best control is the rearing of control and experimental mosquitoes in the same containers throughout their aquatic and adult stages (Habtewold et al., 2017). Here, we achieved this by assaying mosquitoes reared together and were either hemizygous or homozygous positive or negative for the transgene; the latter genotype was used as a reference for the other two. These mosquitoes were produced by crossing the GM lines to the parental G3 line to derive a population of mosquitoes that were hemizygous with respect to the transgene, which was then intercrossed to produce a G2 progeny presenting all three allelic combinations (−/−, +/− and +/+). The G2 mosquitoes were then subjected to the assay, whether transmission blocking or survival, and genotyped for the presence of the modified allele. An added advantage of this protocol is that one can in parallel monitor dose-dependent effects or haplosufficiency of the modified allele, an intrinsic feature of the IGD split drive, where the autonomous drive (here Cas9) and non-autonomous effector (here an AMP) are released separately leading to the effector being initially segregating largely by simple Mendelian genetics. A shortcoming of this otherwise very robust protocol is that all the mosquitoes must be individually genotyped, thus substantially adding to the cost and effort required. Therefore, the protocol requires further streamlining and automation for it to be used in large-scale studies.

The presence and number of oocysts found on the mosquito midgut wall is the most widely used method to assess malaria transmission blocking or reduction caused by an intervention such as a transmission blocking vaccine, drug, or genetic modification. However, this method does not account for any possible defects in oocyst development, a factor critical for computing the parasite extrinsic incubation period (EIP) and thenceforth reproductive rate (R_0_). Examples include the manipulation of the mosquito lipid shuttling machinery that is shown to interfere with oocyst development (Mendes et al., 2008;Rono et al., 2010) and the knockout of the parasite gene *misfit* that leads to oocysts that developmentally arrest but remain detectable and macroscopically indistinguishable from developing oocysts and continue to grow in size (Bushell et al., 2009). Furthermore, recent studies have revealed that the development of oocysts can be significantly delayed or enter dormancy when nutrients acquired through frequent mosquito bloodmeals are scarce and by parasite crowding (Shaw et al., 2020;Habtewold et al., 2021). Thus, supplementing infected mosquitoes with bloodmeals better resembles their natural feeding behavior and physiology and minimize any laboratory biases when evaluating mosquitoes destined for population replacement (Costa et al., 2018;Mota and Mello-Vieira, 2019). In conclusion, a better measure of transmission blockage or reduction is the recording of the presence and abundance of sporozoites in the salivary glands of mosquitoes that are regularly provided with bloodmeals after the initial infectious bloodmeal, a protocol we used in the assays presented here.

Using the protocol described above we find, on the one hand, that Scorpine expression in the CP and AP2 locus does not affect the prevalence and abundance of sporozoites in the mosquito salivary glands 14 days post-infection. This is despite the Sco-CP and ScoG-AP2 lines previously found to harbor more and less oocysts when compared to G3, respectively (Hoermann et al., 2021). These results could be explained by the genetic differences expected to exist between lines, which lead to varied susceptibility to infection and/or the fact that subtle differences in the oocyst load may not translate to differences in sporozoites. The expression of an AMP can affect midgut homeostasis at the time of bloodmeal digestion in ways that are difficult to predict, leading to disturbance of metabolic processes and availability of nutrients for the developing oocyst. Furthermore, quantifying the cumulative abundance of sporozoites in salivary glands as we did here does not report on any possible effects of the intervention on the oocyst developmental rates.

On the other hand, Scorpine expression in the Aper1 locus and docking on the peritrophic matrix leads to a phenotype that merits further investigation with experimental and modeling studies as regards to the potential of this modification to reduce malaria transmission when driven through wild *A*. *gambiae* populations. Aper1-Sco mosquitoes exhibit a significantly shorter lifespan compared to wt, with most of them dying before they become capable of transmitting the parasite. Furthermore, the few GM mosquitoes that survive beyond the parasite EIP show significantly lower prevalence of sporozoite presence, consistent with our previous findings that showed significantly lower infection prevalence and intensity at the oocyst stage (Hoermann et al., 2021). At the same time, GM homozygous and heterozygous mosquitoes appear to harbor significantly more midgut microbiota compared to their homozygous negative siblings 24 hours post-bloodmeal. While Scorpine expression in the midgut may directly compromise blood bolus parasite stages leading to lower infection at both the oocyst and sporozoite stages, the combination of phenotypes we detect could be explained in two additional ways. Firstly, the expression of the exogenous AMP disturbs the midgut homeostasis leading to higher microbiota loads and premature mosquito death. Secondly, fusion of Scorpine with Aper1 compromises the integrity of the type I peritrophic matrix and thus the midgut barrier leading to higher midgut microbiota and systemic infection of the mosquito hemolymph (Rodgers et al., 2017).

Importantly, while self-suppressing, mosquitoes bearing the Aper1-Sco modified allele can effectively compete and mate with wt mosquitoes producing offspring that inherit the modified allele in frequencies comparable to those of the wt allele. Overall, our results suggest that in a field scenario, the reduced survival in conjunction with the reduced sporozoite load of Aper1-Sco GM mosquitoes may significantly impact on parasite transmission in the release area. It would be pertinent, at this stage, to collect additional data from large scale experiments and model the impact of any release of the strain in malaria transmission, especially when combined with a gene drive.

## Data availability statement

The original contributions presented in the study are included in the article. Further inquiries can be directed to the corresponding author.

## Author contributions

Conceptualization, ST, NW and GKC. Methodology, ST, MGI, JC, JP and PC. Formal analysis, ST and GKC. Investigation, ST, JP and GKC. Writing paper, ST and GKC. Supervision, GKC. Project administration, GKC. Funding acquisition, NW and GKC. All authors contributed to the article and approved the submitted version.

## Acknowledgements

The authors thank Temesgen Menberu Kebede for his help with maintaining the mosquito colonies and transgenic lines.

